# Contractility of cardiac and skeletal muscle tissue increases with environmental stiffness

**DOI:** 10.1101/2024.02.23.581737

**Authors:** Delf Kah, Julia Lell, Tina Wach, Marina Spörrer, Claire A. Dessalles, Sandra Wiedenmann, Richard C. Gerum, Silvia L. Vergarajauregui, Tilman U. Esser, David Böhringer, Felix B. Engel, Ingo Thievessen, Ben Fabry

**Affiliations:** Biophysics Group, Department of Physics, Friedrich-Alexander Universität Erlangen-Nürnberg (FAU), Erlangen, Germany; Department of Biochemistry, University of Geneva, 1211 Geneva, Switzerland; Department of Physics and Astronomy, York-University Toronto, Ontario, Canada; Experimental Renal and Cardiovascular Research, Department of Nephropathology, Institute of Pathology, Friedrich-Alexander Universität Erlangen-Nürnberg, Erlangen, Germany

## Abstract

The mechanical interplay between contractility and mechanosensing in striated muscles is of fundamental importance for tissue morphogenesis, load adaptation, and disease progression, but remains poorly understood. In this study, we investigate the dependence of contractile force generation of cardiac and skeletal muscle on environmental stiffness. Using *in vitro* engineered muscle micro-tissues that are attached to flexible elastic pillars, we vary the stiffness of the microenvironment over three orders of magnitude and study its effect on contractility. We find that the active contractile force upon electrical stimulation of both cardiac and skeletal micro-tissues increases with environmental stiffness according to a strong power-law relationship. To explore the role of adhesion-mediated mechanotransduction processes, we deplete the focal adhesion protein β-parvin in skeletal micro-tissues. This reduces the absolute contractile force but leaves the mechanoresponsiveness unaffected. Our findings highlight the influence of external stiffness on the adaptive behavior of muscle tissue and shed light on the complex mechanoadaptation processes in striated muscle.

## Introduction

The mechanical properties of the microenvironment are critical for cell and tissue morphogenesis, homeostasis, regeneration, and disease progression [1, 2]. Striated muscle in particular is well known to respond sensitively to the mechanical environment [3, 4]. For example, the heart adapts its stroke volume to the hemodynamic load according to the Frank-Starling mechanism [5], and shows hypertrophic responses to long-term increases in load [6, 7]. Similarly, skeletal muscles increase their maximum contractile force by long-term training under isometric conditions [8], but develop atrophy under reduced mechanical load *e.g.* under microgravity conditions in space [9].

During embryonic development, striated muscle tissue forms preferentially along mechanical stiffness gradients [10]. A related effect can also be observed *in vitro*: micro-tissues that are formed between two flexible cantilevers increase their contractile force with increasing cantilever stiffness [10, 11]. The molecular basis of the underlying mechanosensing and mechanochemical transduction processes, however, are still poorly understood. This is partly due to the difficulty to impose defined loading conditions to muscle cells and tissue grown in traditional two-dimensional (2-D) cell culture systems. Moreover, 2-D muscle cannot replicate the three-dimensional (3-D) adhesive microenvironment of native muscle tissue with its specialized cellular mechanosensing structures such as costameres and intercalated disks [12, 13].

The distinctive feature of striated muscle compared to other cell types is the ability to actively contract in response to electrical stimulation. At the molecular level, muscle contraction results from the interaction of sarcomeric actin and myosin. The sarcomeres are laterally coupled to the extracellular matrix (ECM) via costameres. The transmembrane force coupling to the ECM is established in large part through integrins. Intracellularly, integrins form an adhesion complex consisting of talin, vinculin, and numerous other focal adhesion proteins that in turn connect to the cytoskeleton. Most of the focal adhesion proteins, including integrins themselves, have been implicated in mechanochemical signaling [14, 15]. Recently, we have shown that a critical mechanoresponsive signaling hub in cardiac muscle is the focal adhesion protein β-parvin [16]. In neonatal rat ventricular cardiomyocytes, β-parvin connects to integrins via integrin-linked kinase and activates Rac1 in response to mechanical load, thereby controlling cell shape, sarcomere assembly, and contractility. Since β-parvin is also highly expressed in skeletal muscle tissue, it is reasonable to assume that it may have mechanoregulatory properties in skeletal muscle similar to those in cardiac muscle.

Here, we use *in vitro* engineered muscle micro-tissues of cardiac or skeletal origin to study how the developing muscle responds to the external stiffness of the microenvironment. In our study, muscle micro-tissues are formed between two elastic polydimethylsiloxane (PDMS) pillars with an effective spring constant that can be adjusted over three orders of magnitude. We find that the active contractile forces of cardiac and skeletal muscle micro-tissues strongly respond to the spring constant of the pillars. In particular, the electrically-triggered contractile force increases with higher pillar stiffness according to a power-law relationship, with power-law exponents between 0.3 and 1.0 depending on cell type and time in culture. We furthermore show that this mechanoadaptation is independent of absolute contractility, as skeletal micro-tissues with a β-parvin knockdown exhibit dramatically reduced active and static forces and contraction velocity compared to control siRNA micro-tissues, but still show a similar power-law relationship between active force and environmental stiffness.

## Results

### Micro-tissues as a model for structural and functional maturation of skeletal muscle

To generate muscle micro-tissues, we added 5000 C2C12 skeletal muscle cells suspended in 6 µl of unpolymerized collagen-I/Matrigel solution to polydimethylsiloxane (PDMS) chambers containing two pillars (0.5 mm diameter, placed 2 mm apart) (Fig. 1A). Upon matrix polymerization, the cells adhered to the matrix fibers and exerted contractile forces, resulting in matrix remodeling and the formation of a micro-tissue within 24 h (Fig. 1B, S1). The height at which the micro-tissues attach between the pillars was tuned by forming a first layer of cell-free matrix at the bottom of the PDMS chamber with a defined volume and allowing it to polymerize for one hour prior to cell seeding in a second collagen layer on top of the bottom layer. After 24 hours, we replaced the culture medium with a differentiation medium, supplemented with 0.5% horse serum and 0.5% ITS (insulin, transferrin, selenium), to initiate myocyte differentiation.

**Fig. 1.**
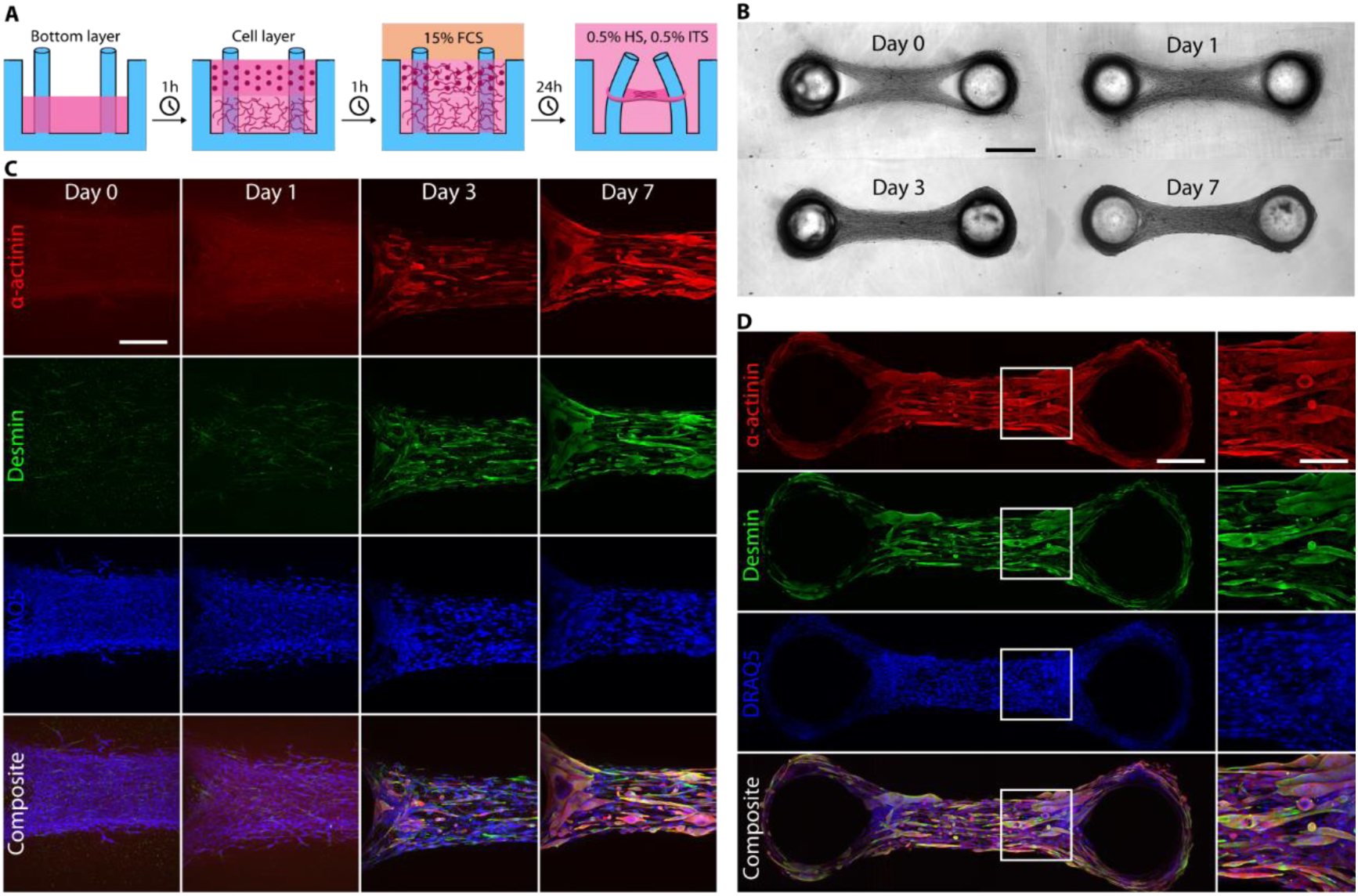
Formation and maturation of micro-tissues generated from C2C12 cells. **(A),** Schematic of the experimental protocol for micro-tissues fabrication. In a two-step process, we pipette a layer of cells mixed in an unpolymerized collagen/Matrigel matrix into the PDMS well on top of a prepolymerized bottom layer without cells. Within 24 hours, micro-tissues form in a cell culture medium containing 15% FCS, and we subsequently switch to a differentiation medium containing 0.5% HS and 0.5% ITS for the remainder of the experiment. **(B),** Bright-field images of a micro-tissue over several days in culture. Scale bar: 500 µm. **(C),** Maximum intensity projections of confocal image stacks of micro-tissues fixed at 0, 1, 3, and 7 days after switching to differentiation medium. Tissues are stained for α-actinin, desmin, and nuclei (DRAQ5). Excitation intensities and detector gains were the same for all samples. The gradual increase in α-actinin and desmin intensities over time demonstrate maturation of the myocytes in culture. Scale bar: 200 µm. **(D),** Maximum intensity projections of confocal image stacks over the entire length of a micro-tissue (fixed on day 7) stained as before. The close-up images show parallel orientation of the myocytes along the axis connecting the pillars. Scale bar: 200 µm. Scale bar in close-up: 100 µm.

Over the following days, the myocytes started to exert measurable forces on the pillars, causing further matrix compaction and fiber alignment between the pillars (Fig. 1A, B). Over the course of seven days of differentiation, the myocytes elongated and aligned between the pillars. In addition, they showed increasing expression levels of the muscle differentiation markers α-actinin and the intermediate filament desmin, as revealed by immunofluorescence staining (Fig. 1C, D).

Already one day after cell seeding, the collective contractile forces exerted by the myocytes caused the micro-tissues to pull on the pillars, resulting in a pillar deflection that was visible with bright-field imaging. After 4 days of differentiation culture, the muscle tissues started to respond to electrical pacing, as seen by a measurable pillar deflection with every pulse. The force acting on the pillars was then calculated according to Hooke’s law (Eq. 1) from the pillar deflection (Fig. 2A,B) multiplied by the pillar spring constant (Fig. 2C). We refer to the continuous baseline force as the *static force*, and we refer to the additional force above the static force that is observed in response to electrical stimulation as the *active force* (Fig. 2D).

**Fig. 2.**
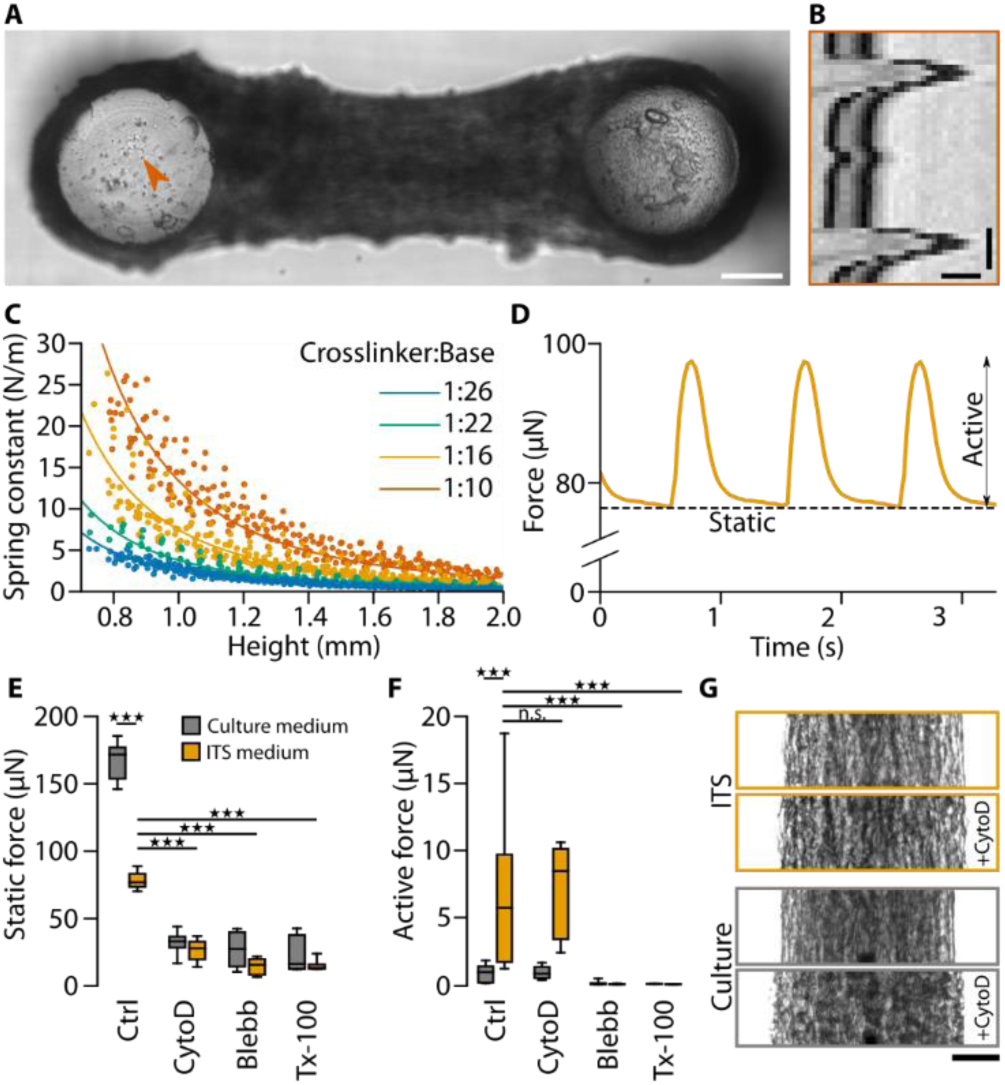
Contractile forces in C2C12 micro-tissues. **(A),** Bright-field image of a C2C12 micro-tissue focused at the top of the PDMS pillars. The orange arrowhead indicates a high-contrast point that is used for particle image velocimetry analysis. Scale bar: 200 µm. **(B),** Kymographic representation of the movement of the point indicated in (A) in response to active tissue contraction. Scale bars: 10 µm and 250 ms. **(C),** Calibration curves to determine the spring constants of the PDMS pillar. The spring constants were measured from the force-deflection relationship of pillars at different heights and for PDMS with different crosslinker-to-base ratios. Dots represent slopes of individual force-deflection measurements, and lines represent fits to the data points using the Euler-Bernoulli beam equation. **(D),** Static and active contractility of the micro-tissue shown in (A). The stimulation frequency was 1 Hz. **(E),** Static contractile forces of C2C12 micro-tissues after 7 days in culture. One group of micro-tissues was prepared as described in the text (*i.e.*, using ITS differentiation medium), and one group was kept in culture medium for the entire 7 days (instead of switching to ITS medium after 24 hours) so that the myocytes did not differentiate. We measured the static contractile forces under control conditions (Ctrl; Dimethylsulfoxid at a concentration of 1:100), after adding either 10 µM cytochalasin-D (cyto-D), or 100 µM blebbistatin (Blebb) to the medium for 1 hour, or after adding Triton X-100 (Tx-100) to the medium for 10 minutes; n.s., P≥0.05; ★★★, P<0.001. **(F),** Same as (E) for active contractile forces. **(G),** Contrast enhanced bright-field images of exemplary micro-tissues before and after treatment with cyto-D. One micro-tissue was cultured in ITS medium and the other in culture medium before drug treatment. Scale bar: 100 µm.

The static force of C2C12 micro-tissues increased over time in differentiation culture. In PDMS devices of intermediate stiffness (average pillar spring constant ∼ 6 N/m), the static force reached 76 µN on average after one week, and the active force was 7.5 µN (Fig. 2E-F). In contrast, undifferentiated C2C12 micro-tissues that were continuously exposed to culture medium but not differentiation medium showed higher static forces (166 µN on average), but almost no measurable active forces (0.9 µN on average), suggesting that the static force is mainly exerted by undifferentiated C2C12 cells, whereas the active force is mainly exerted by differentiated muscle cells.

To further explore the origins of static and active contractility, we measured forces after treatment with inhibitors for different components of the contractile machinery (Fig. 2E,F). Addition of 100 µM of the myosin II inhibitor blebbistatin for one hour completely impaired active contractility in all micro-tissues (active forces were less than 0.1 µN, resulting solely from noise in the pillar deflection measurements). This confirms that myosin is essential for muscle contractility. Cell permeabilization with Triton X-100 at a concentration of 0.5% for ten minutes also completely impaired active contractility, confirming that an intact sarcolemma is essential for initiating contractions through electrical stimulation. Similarly, static forces were strongly reduced and fell to 27 µN in undifferentiated and 14 µN in differentiated micro-tissues upon blebbistatin-treatment, and to 24 µN in undifferentiated and 16 µN in differentiated micro-tissues upon Triton X-100 treatment. These findings are consistent with the requirement of myosin II activity for contractile force generation in non-muscle cells [17].

Since sarcomeric actin filaments are stably capped at their barbed ends in the Z-discs of myofibers [18, 19], we hypothesized that the active forces in differentiated C2C12 muscle-micro-tissues are less susceptible to barbed-end depolymerization than the static forces. To test this, we treated differentiated and undifferentiated micro-tissues with high doses of the F-actin barbed-end depolymerizing fungal toxin cytochalasin D (cyto-D). Treatment with 10 µM cyto-D for one hour reduced the static forces in differentiated and undifferentiated micro-tissues to similar levels as seen after treatment with blebbistatin or Triton-X (26 µN in differentiated and 32 µN in undifferentiated micro-tissues). By contrast, cyto-D treatment had no influence on active contractile forces in differentiated micro-tissues (7.1 µN). In addition, bright-field imaging revealed that after treatment with cyto-D, undifferentiated micro-tissues increased in width, and many cells appeared rounded (Fig. 2G), whereas tissue widening and cell rounding were less pronounced in differentiated micro-tissues. The observation that cyto-D has an inhibitory effect only on static but not on active forces strongly supports our hypothesis that active forces are generated predominantly by differentiated cells where the barbed ends of sarcomeric actin filaments are stably capped, rendering the cells insensitive to cyto-D treatment, while cyto-D can initiate barbed-end depolymerization of the non-sarcomeric F-actin in undifferentiated myocyte precursors, resulting in reduced static forces. The presence of appreciable static forces after treatments with cyto-D, blebbistatin and Triton X-100 furthermore suggests that a part of the static force originates from stored mechanical tension within remodeled collagen fibers.

Next, we sought to understand the role of the ECM in the maturation of the micro-tissues. The differentiation and adhesion of myoblasts and myotubes to the ECM requires the engagement of adhesion receptors such as integrins or the dystrophin-glycoprotein complex to laminin, a critical adhesive component of the extracellular matrix surrounding skeletal myofibers and cardiac myocytes [20, 21]. Since laminin is a major component of Matrigel [22], we tested whether the presence of Matrigel was required for proper C2C12-micro-tissue formation and contraction. C2C12 cells mixed into pure collagen-I (COL), without the addition of Matrigel, did not develop uniformly compact micro-tissues but only formed loose tissues interspersed with denser aggregates over the course of 7 days (Fig. 3A). Importantly, these micro-tissues were unable to generate active contractile forces (Fig. 3B). These results suggest that laminin promoted C2C12-myoblast differentiation in our micro-tissues, in line with our previous results that active contractility requires differentiated cells.

**Fig. 3.**
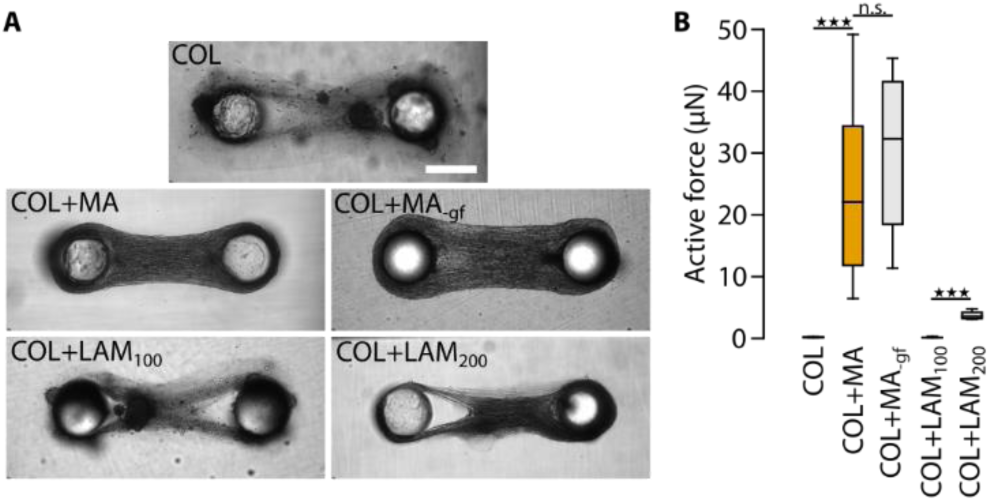
Laminin is the critical component for C2C12 micro-tissue compaction and maturation. **(A),** Bright-field images of representative micro-tissues for all conditions. Scale bar: 500 µm. **(B),** Active contractile forces of C2C12 micro-tissues prepared with ECMs of different compositions: collagen only (COL), collagen and 1 mg/ml Matrigel (COL+MA), collagen and 1 mg/ml growth factor-reduced Matrigel (COL+MA_-gf_), collagen and 100 µg/ml laminin (COL+LAM_100_), and collagen and 200 µg/ml laminin (COL+LAM_200_); n.s., P≥0.05; ★★★, P<0.001.

To test whether the impaired muscle differentiation in the absence of Matrigel was indeed due to a lack of laminin, or due to the lack of growth factors that are also part of Matrigel, we compared C2C12 micro-tissues with collagen-I containing 1 mg/ml of either regular (COL+MA) or growth factor-reduced Matrigel (COL+MA_-gf_), or 100 µg/ml (COL+LAM_100_), or 200 µg/ml (COL+LAM_200_) pure laminin isolated from Engelbreth-Holm-Swarm murine sarcoma basement membrane. Micro-tissues formed in the presence of growth factor-reduced Matrigel resembled control COL+MA tissues in terms of morphology and contractility (COL+MA = 25.0 µN; COL+MA_-gf_ =29.9 µN; two-tailed t-test: P=0.42). Similar to pure COL micro-tissues, COL+LAM_100_ micro-tissues showed a low degree of compaction, presence of aggregates, and no active forces. COL+LAM_200_ micro-tissues, however, compacted better and developed low levels of active contractile forces (mean: 3.8 µN), indicating that even moderate levels of pure laminin were able to improve micro-tissue formation and contraction. These data show that lamin 1 is important for myotube differentiation, while matrix-bound growth factors present in Matrigel are not. However, addition of 200 µg/ml laminin-1 was not sufficient to restore the full contractility seen with the addition of Matrigel, suggesting that the high amount of laminin in Matrigel (approx. 600 µg/ml, [22]) was required for full contractility or that other Matrigel components also contributed to myotube differentiation.

Together, these data show that C2C12-based micro-tissues recapitulate key features of skeletal muscle, *i.e.* myoblast elongation and differentiation, electrical excitability and contractile force generation, suggesting that they can be used as functional *in vitro* model for striated muscle tissue.

### Neonatal rat cardiomyocytes form micro-tissues with intact electrical coupling

Similar to skeletal muscle, cardiac muscle also is a mechanically highly loaded tissue with a complex cell-adhesive microenvironment that cannot be adequately replicated using 2-D cell culture systems. Moreover, neonatal rat ventricular cardiomyocytes (NRVC), the most commonly used cardiomyocyte model system, have not yet undergone postnatal terminal differentiation, lacking key features of adult cardiomyocytes such as a fully organized sarcomere apparatus and a rod-like cell shape. Hence, we next tested whether NRVCs were able to form micro-tissues in our system and whether cells and micro-tissues show features of adult cardiomyocytes and myocardial tissue, respectively. Over three days in culture, NRVCs had fully compacted the ECM into micro-tissues (Fig. 4A). NRVCs best formed micro-tissues using a lower collagen concentration (0.3 mg/ml) compared to C2C12 cells (0.6 mg/ml). Tissue formation did not require the addition of Matrigel, possibly because NRVCs were already partly differentiated *in vivo*. Notably, confocal imaging of α-actinin revealed an elongated morphology of NRVCs and a homogenous cross-striation pattern reminiscent of adult myocardial tissue *in vivo* (Fig. 4B). In addition, NRVC micro-tissues displayed spontaneous, as well as electrically induced active contractions already three days after cell seeding. The static forces of NRVC micro-tissues were one order of magnitude lower than those of C2C12 micro-tissues (typically 1-10 µN), while the active forces were comparable (typically 20 µN). The active contraction of NRVC micro-tissues followed the frequency of the electrical stimulation up to about 4 Hz (Fig. 4C). At higher frequencies, the micro-tissues did not fully relax between pacing pulses, but single force twitches did also not fuse to a tetanic force as seen in skeletal muscle micro-tissues [23] and still showed oscillations in amplitude. Consequently, the maximum active forces at higher stimulation frequencies (2, 4, 8, 10 Hz) remained unchanged compared to single pulse stimulation (0.5 and 1 Hz). Single twitches were still clearly observable even at high pulsing frequencies (10 Hz).

**Fig. 4.**
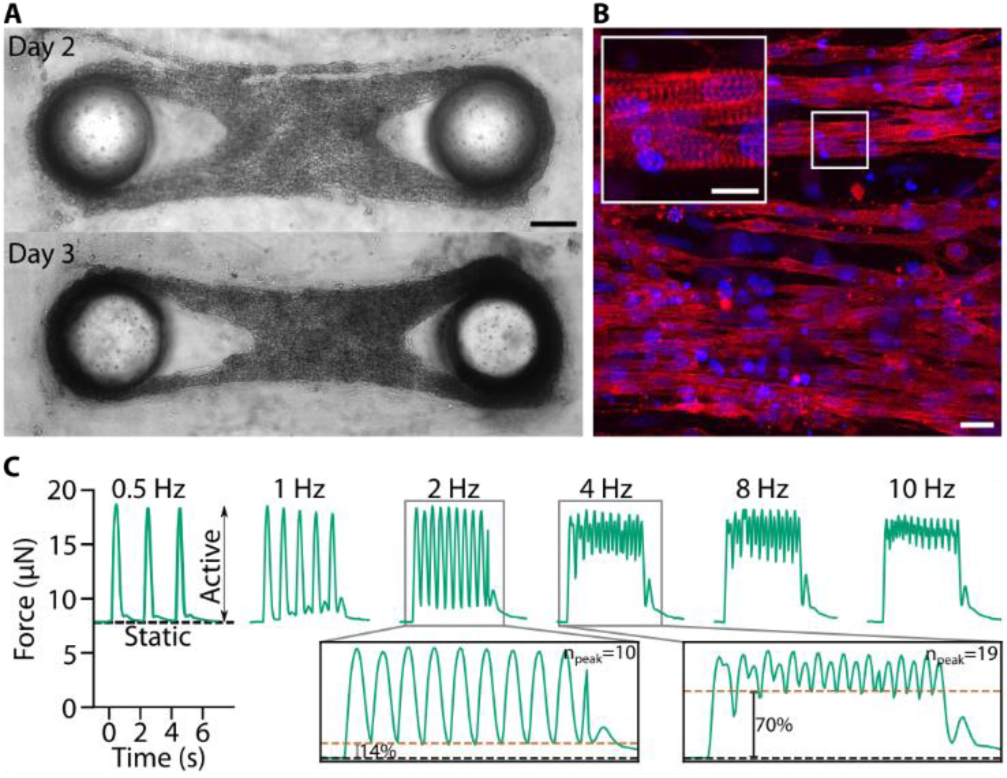
Micro-tissues fabricated from neonatal rat cardiomyocytes. **(A),** Bright-field images of a representative micro-tissue on day 2 and 3 after cell seeding. Scale bar: 200 µm. **(B),** Maximum projections of confocal image stacks of a micro-tissue fixed 5 days after cell seeding and stained for α-actinin (red) and nuclei (DRAQ5, blue). Scale bar: 20 µm; scale bar inset: 10 µm. **(C),** Static and active contractility of a cardiac micro-tissue at different stimulation frequencies. In the insets, the black dashed lines represent the static forces and the orange dashed lines represent the minimum total force between pacing pulses. Percentages indicate the average minimum active force between pacing pulses, relative to the average maximum active force.

These data show that NRVC-based micro-tissues replicate essential features of terminally differentiated myocardial tissue, such as cardiomyocyte elongation and cross-striation as well as electrically inducible contraction, suggesting that, similar to our C2C12 micro-tissues, they are well suited for *in vitro* studies of cardiac muscle tissue.

### The active contractility of skeletal and cardiac muscle micro-tissues is mechanoadaptive

To investigate cellular and molecular mechanisms of striated muscle mechanoresponses, we next made use of a key feature of our system ‒ the ability to tune the environmental stiffness to which the micro-tissues are exposed. The relevant measure of environmental stiffness is the effective spring constant *k* of the PDMS pillars at the height *h* of the micro-tissue attachment (with respect to the bottom of the PDMS chamber). As predicted by Euler-Bernoulli beam theory, the relationship between stiffness and micro-tissue height is *k* ∼ *E* × ℎ^−3^, where *E* is the Young’s modulus of the PDMS (Fig. 2C). Consequently, there are two ways to tune the environmental stiffness: i) by varying the volume of the bottom layer during the fabrication process so that the micro-tissues form at different heights *h* between the pillars (Fig. 5A), and ii) by using PDMS devices with different Young’s modulus *E*, *i.e.* with different crosslinker-to-base ratios.

**Fig. 5.**
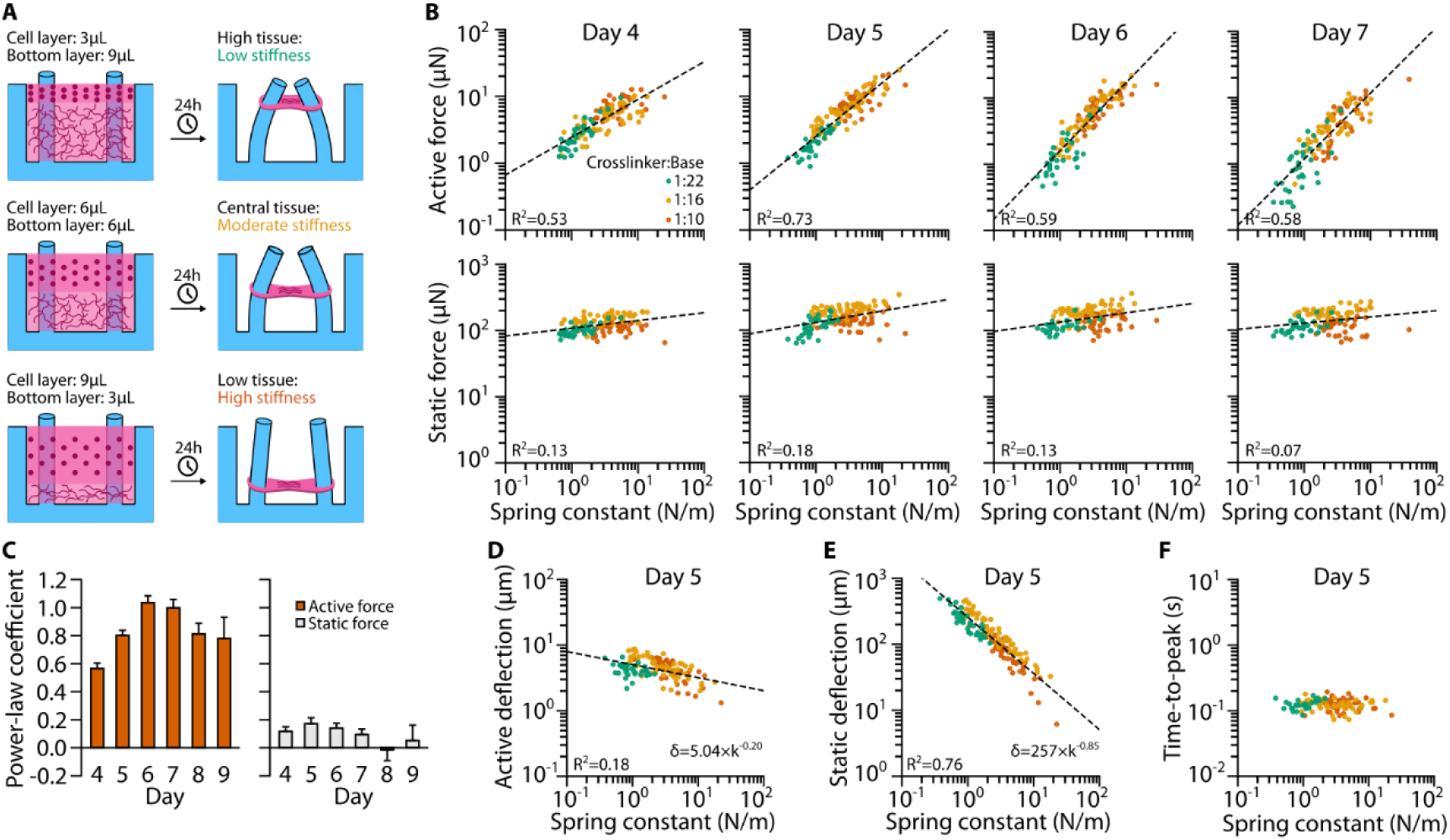
Mechanoadaptation of C2C12 micro-tissues. **(A),** Schematic of the relationship between bottom layer volume and tissue height. After 24 hours of initial compaction, the micro-tissues are approximately attached to the PDMS pillars at the level of the bottom/cell layer interface. Since the effective spring constant *k* of the pillars depends on the micro-tissue height, a high/intermediate/low bottom layer (9/6/3 µl) will result in a low/moderate/high environmental stiffness. **(B),** Scatter plots of active and static forces of C2C12 micro-tissues versus *k* measured on days 4-7 after initialization of differentiation. Dashed lines represent power-law fits. **(C),** Power-law coefficients for the same fits (with days 8&9 added). Shown are means and SDs derived from bootstrapping. **(D),** Scatter plot of active deflections over *k* for day 5 with power-law fit. **(E),** Same as (D) but for static deflection. **(F),** Scatter plot of times-to-peak (*i.e.*, the time at which the micro-tissues reach maximum deflection after electrical stimulation) versus *k* at day 5.

To investigate the influence of the mechanical environment and of culture time on the contractile performance, we first fabricated C2C12 micro-tissues with different *E* and *h* and measured the active and static contractile forces every day from day 4-9 after switching to differentiation medium (Fig. 5B). We found that the active forces *F* increased with environmental stiffness *k*, and that the relationship can be well approximated by a simple power-law relationship of the form *F*(*k*) = *a* × (*k*/*k*_0_)^*b*^, where the factor *a* is a measure of contractile force at a nominal spring constant of *k_0_* = 1 N/m, and the power-law coefficient *b* is a measure of mechanoadaptation (Fig. 5C). A power-law exponent of zero corresponds to a tissue that does not adapt to environmental stiffness, and an exponent of unity corresponds to a tissue that increases its contractile force linearly with increasing environmental stiffness.

On day 4 ‒ the first day with measurable, active contraction ‒ active forces increased with *k* according to a power-law exponent *b*=0.57±0.04 (error derived by bootstrapping; *R^2^*=0.53), *i.e.* roughly with the square root of environmental stiffness (Fig. 5B). Over the next two days in culture, the degree of mechanoadaptation increased further, reaching a maximum power-law coefficient of 1.03±0.05 on day 6 (*R^2^*=0.59), hence contractility increased nearly linearly with environmental stiffness. By contrast, the static contractile force increased only weakly with *k* and reached a maximum power-law coefficient of 0.17±0.04 (*R^2^*=0.18) on day 5 (Fig. 5B). These data demonstrate a pronounced mechanoresponse of C2C12 micro-tissues and furthermore suggest that this mechanoadaptation is caused by differentiated myotubes, but not myoblasts.

We next explored how muscle shortening and hence pillar deflection changed with the spring constant, since the active force of striated muscle may decrease with increasing amount of shortening as the internal resistance against filament sliding increases [24]. To this end, we correlated pillar deflection for active and static forces with the corresponding spring constant. Consistent with the absence of a clear mechanoresponse of static force generation, the pillar deflection induced by static forces decreased over several orders of magnitude with increasing pillar stiffness. By contrast, the pillar deflection due to active forces was nearly constant, regardless of pillar stiffness. For example, on day 5, the static pillar deflection was strongly correlated with *k* (*R^2^*=0.76) and had a power-law coefficient of −0.85 (Fig. 5E), while the active pillar deflection was only weakly correlated with *k* (*R^2^*=0.18) and had a power-law coefficient of −0.20 (Fig. 5D). Together with the observation that the maximum amount of shortening was only 2% of the total tissue length, this finding suggests that the active forces of C2C12 micro-tissues in a softer microenvironment are not limited by filament sliding constraints or by a limited capacity to shorten. Consistent with this, the time required for the micro-tissues to reach maximum active contraction (time-to-peak) was independent of the spring constant (Fig. 5F).

We next tested whether cardiomyocyte micro-tissues were also adapting to pillar stiffness and observed a less pronounced but still strong mechanoadaptation response in micro-tissues prepared from NRVCs (Fig. 6A,B). These micro-tissues first exhibited active contraction at day 3 after initial cell seeding, with a power-law coefficient of the active forces versus *k* relationship of 0.48±0.04 (*R^2^*=0.59). Thereafter, the degree of mechanoadaptation decreased slightly with culture time, and the power-law coefficient was between 0.35-0.39 on days 4-6 and decreased to 0.25 on day 7. These data demonstrate that the mechanoadaptation is not limited to skeletal muscle micro-tissues, suggesting that the underlying mechanisms may be general to different types of striated muscle tissues.

**Fig. 6.**
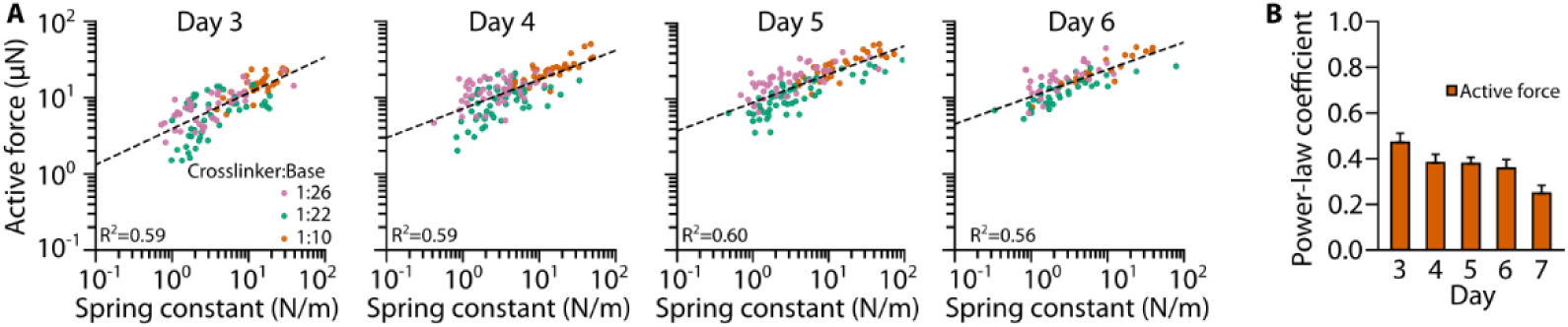
Mechanoadaptation of NRVC micro-tissues. **(A),** Scatter plots of NRVC micro-tissue active forces versus *k* measured on days 3-6 after cell seeding. Dashed lines represent power-law fits. **(B),** Power-law coefficients for the same fits (with day 7 added). Shown are means and SDs derived from bootstrapping.

### Only contractile force but not stiffness of muscle micro-tissues is mechanoadaptive

Our PDMS devices are designed such that they can be mounted on a stepper motor-driven uniaxial cell stretcher device [25]. This allows us to determine the stiffness of micro-tissues by applying an external strain and evaluating the resulting tissue force and tissue strain. For this purpose, we pre-stretched PDMS devices by 15% and generated C2C12 micro-tissues as described above, while maintaining the pre-stretch throughout the culture period. On the day of measurement, we mounted the PDMS devices on the cell stretcher and gradually reduced the amount of stretch to zero (Fig. 7A). We chose this approach instead of stretching unstretched samples because micro-tissues are also shortening during active and static contraction.

**Fig. 7.**
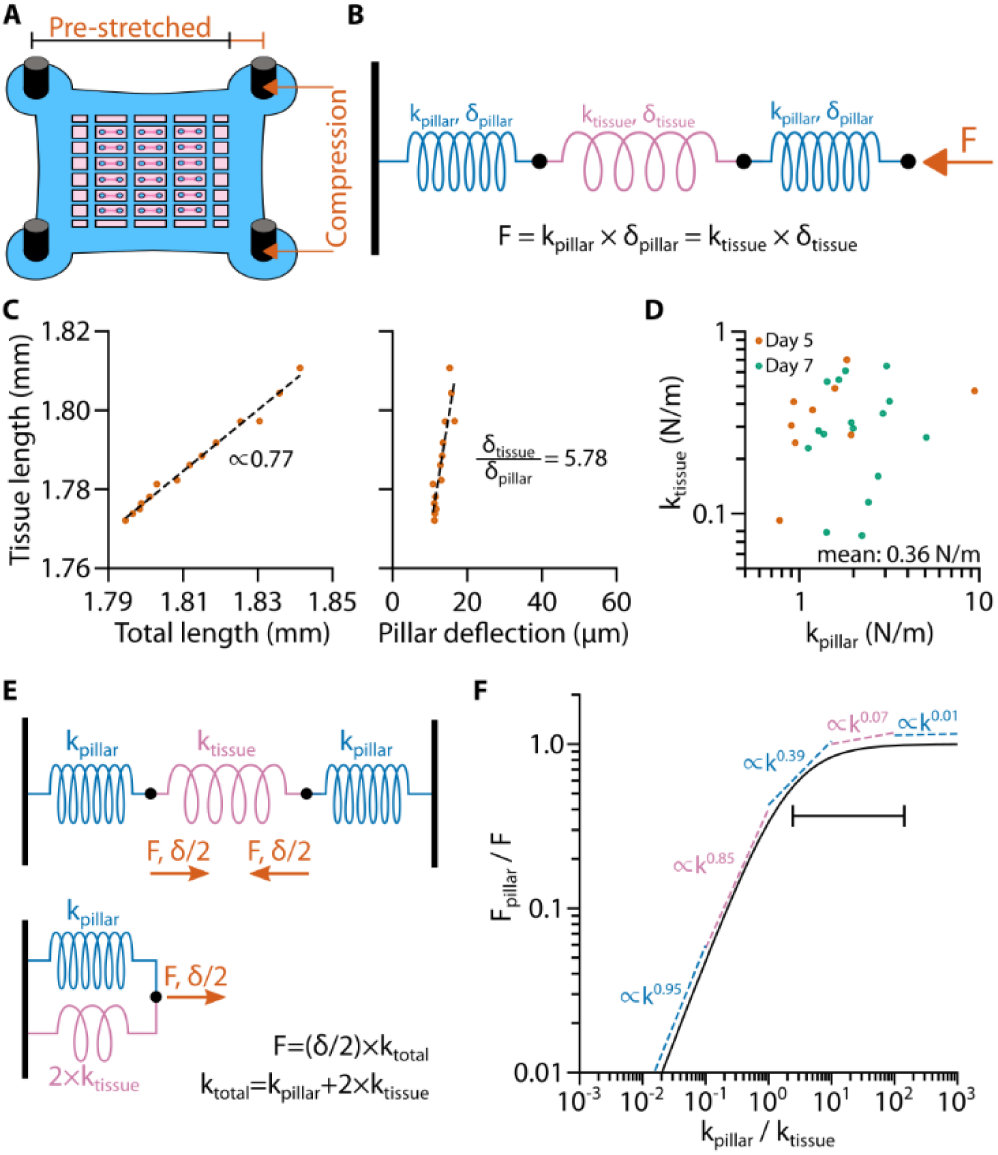
Spring models for contractility and measurement of micro-tissue stiffness. **(A),** Schematic of a pre-stretched PDMS device. **(B),** Three-spring-model of a micro-tissues between two pillars during stretch/compression. Note that during uniaxial stretch/compression, one end of the serial spring assembly is fixed while the external force acts on the other end. **(C),** Scatter plot of tissue length versus total length (tissue length + pillar deflections) and tissue length versus deflection of one pillar. Dashed Lines represent linear fits. **(D),** Scatter plot of the calculated values of *k_tissu_*_e_ versus the corresponding values of *k_pillar_*. **(E),** Three-springs-model of a micro-tissues between two pillars during active/static contraction and equivalent circuit of one half of this arrangement (from one wall to the center of the micro-tissues). **(F),** Line plot of the analytical expression *F_pillar_* / *F* versus *k_pillar_ / k_tissue_*. Dashed lines represent power-law fits of *F_pillar_* / *F* in different decades of *k_pillar_ / k_tissue_*. The line with vertical endpoints represents the stiffness range of the PDMS pillars used in previous experiments.

The relationship between the external force *F*, tissue shortening *δ_tissue_*, and pillar deflection *δ_pillar_* can be modeled by a serial arrangement of three springs (Fig. 7B): a spring with spring constant *k_tissue_*, representing the micro-tissue, is connected at both ends to a spring with spring constant *k_pillar_*, representing the pillars. In this model, *F* acts on the springs at one end (representing the moving end of the cell stretcher), while the other end is attached to a wall (representing the stationary end of the cell stretcher). Due to their serial arrangement, *F* acts equally on each individual spring. The stepwise compression of the PDMS device allows us to effectively vary *F* and measure the absolute micro-tissue length and the absolute pillar deflections via bright-field imaging.

The ratio of the changes in micro-tissue length with respect to the total length or with respect to the deflection of a single pillar can then be used to fit the ratio *δ_tissue_*/*δ_pillar_* (Fig. 7C). Since we can determine *k_pillar_* from calibration measurements as described above, this method allows us to estimate *k_tissue_*.

We performed these experiments with C2C12 micro-tissues for varying pillar stiffness on days 4 and 6 after switching to differentiation medium (the days on which we had measured the lowest and highest degree of mechanoadaptation, respectively). We saw on both days that *k_tissue_* was independent of *k_pillar_*, demonstrating that the stiffness of the micro-tissues is not mechanoadaptive. Furthermore, *k_tissue_* was not significantly different between days (two-tailed t-test: P=0.750), with an average spring constant of all measured micro-tissues of 0.36 ± 0.04 N/m (mean±SE; Fig. 7D), corresponding to a Young’s modulus of 9.2 kPa (for an effective tissue radius of 150 µm).

The effective spring constant of a C2C12 micro-tissue is in the same order of magnitude as the spring constant of the softest PDMS pillars used in this study. This creates a potential source of misinterpretation of mechanoadaptation responses, as the deflection of a soft pillar (which corresponds to the shortening of the tissue) may be so large that the internal tissue stiffness causes a substantial counter-force upon tissue compression that limits further shortening. To distinguish mechanoadaptation of the tissue from such passive force limitation, we calculated how tissue stiffness influences the relationship between contractile force and environmental stiffness for a tissue that is not mechanoadaptive (Fig. 7E,F, see Methods). Accordingly, for a non-mechanoadaptive tissue generating a contractile force *F*, the force measured from the pillar deflection (*F_pillar_*) increases approximately linearly with environmental stiffness *k_pillar_* if *k_tissue_* ≫ *k_pillar_*. For *k_tissue_* ≪ *k_pillar_*, the measured *F_pillar_* approaches the true tissue force *F*.

In most of our experiments, the pillar stiffness is considerably higher than the tissue stiffness, ranging from *2×k_tissue_* to *150×k_tissue_*, as indicated by the bar in Fig. 7F. In this region, we see only a weak dependency between pillar force and tissue stiffness for non-mechanoadaptive model tissues, with power-law exponents on the order of 0.1 and lower. Therefore, we conclude that the power-law dependency of active forces with increasing pillar stiffness with values between 0.3 and 1.0 that we see in our measurements is a signature of true mechanoadaptation, even on days where the power-law exponent was lowest (day 4 for skeletal muscle tissues, day 7 for cardiac muscle tissues) (Fig. S2).

### Knockdown of the focal adhesion protein β-parvin impairs contractile forces, but not mechanoadaptation of C2C12 micro-tissues

The only way for the micro-tissues to probe the stiffness of their environment is to exert force and somehow quantify the resulting length change. We therefore investigated whether the mechanoadaptation of micro-tissues is affected by the maximum force magnitude that they can generate. Previously, we have identified a role of the focal adhesion protein β-parvin in regulating myocyte shape, sarcomere assembly, and contractility in cardiac cells in response to mechanical load [16]. Micro-tissues prepared from Parvb-siRNA-treated NRVCs exhibited reduced active forces compared to micro-tissues prepared from control-siRNA-treated NRVCs [16]. Since β-parvin is also highly expressed in skeletal muscle [16], we tested whether β-parvin knockdown affected the contractility and mechanoadaptation of C2C12 micro-tissues.

To this end, we fabricated C2C12 micro-tissues between PDMS pillars with different spring constants as described above, using cells treated with either control-(siControl) or Parvb-(siParvb) siRNA 24 hours prior to tissue formation. SiControl micro-tissues were phenotypically similar to untreated C2C12 micro-tissues (Fig. 8A) and started to actively contract on day 4. By contrast, siParvb micro-tissues were not able to fully compact the collagen matrix. Consistent with our observations in NRVC micro-tissues [16], absolute active forces were strongly reduced by β-parvin knockdown on day 5, but recovered to the level of siControl micro-tissues by day 7 (Fig. 8B,C) likely due to siRNA-turnover or siRNA-dilution caused by cell division over time. The static forces were also significantly decreased by β-parvin knockdown but did not recover to the levels of siControl micro-tissues after day 7. Note that the distribution of pillar spring constants in siParvb and siControl micro-tissues was comparable (Fig. S3).

**Fig. 8.**
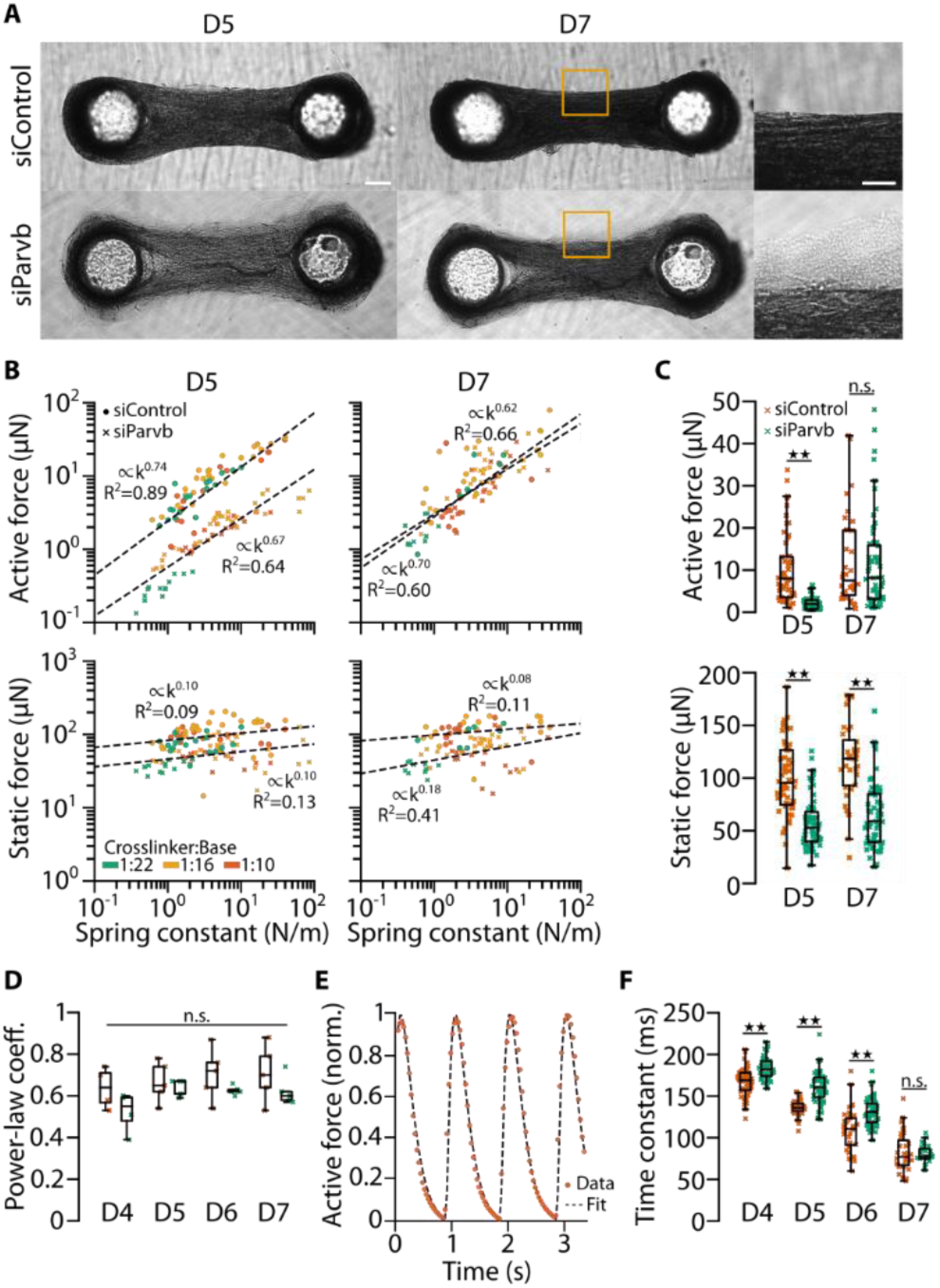
Knockdown of β-parvin impairs absolute contractile forces, but not mechanoadaptation in C2C12 micro-tissues. **(A),** Bright-field images of micro-tissues prepared from control- and Parvb-siRNA-treated C2C12 cells. Images were taken five and seven days (D5 and D7) after switching to differentiation medium. Scale bar: 200 µm. Scale bar in close-up: 100 µm. **(B),** Scatter plots of active and static forces of control-siRNA and knockdown micro-tissues at D5 and D7. Dashed lines represent power-law fits. **(C),** Bar plots of the same data. Bootstrapping; n.s., P≥0.05; ★★, P<0.01. **(D),** Bar plots of the power-law coefficients of fits of active forces and spring constants from 4-5 independent experiments on D5-7. Two-tailed t-test; n.s., P≥0.05. **(E),** Example time course of normalized active contractile force of a control siRNA-treated micro-tissue (dots) fitted by a twofold low-pass filtered Delta-Dirac peak (dotted line) at the time point when the force exhibits local minima. **(F),** Bar plots of the time constants of low-pass filtered active contractions control- and Parvb-siRNA-treated micro-tissues at D5-7. Two-tailed t-test; n.s., P≥0.05; ★★, P<0.01.

Strikingly, the mechanoadaptation of the active forces was almost unaffected in siParvb micro-tissues at day 5 and 7 (Fig. 8B,D). The average power-law coefficient of the active force versus spring constant relationship for siControl micro-tissues was slightly but not significantly higher than that of siParvb micro-tissues for all days (siControl: 0.64-0.71; siParvb: 0.52-0.64, n = 5 independent experiments each with >100 tissues). This indicates that micro-tissues remain fully mechanoresponsive even when the active and static forces are markedly reduced. A plausible explanation is that the expression levels of β-parvin after transient siRNA knock-down recover inhomogeneously over time across the cell population, so that the active tissue force, although it is reduced, is generated by differentiated myotubes that may be fewer in number but are still fully mechanoresponsive.

To test the idea that a temporary β-parvin knockdown delayed muscle differentiation in C2C12 micro-tissues, we analyzed the time course of micro-tissue contraction. The active force response after a single pulse (twitch force) can be modeled as a Dirac-pulse that is low-pass filtered twice with the same time constant *τ*, as explained in Methods.

We find that *τ* of control and β-parvin knockdown tissues decreased over time after initiating muscle differentiation, consistent with the notion that *τ* reports on the degree of muscle differentiation (Fig. 8F). The twitch time constant was significantly longer in siParvb micro-tissues compared to siControl micro-tissues (day 5; siControl: 137 ms; siParvb: 161 ms; two-tailed t-test: P<10^-10^), but converged to similar levels over time (day 7; siControl: 82 ms; siParvb: 81 ms; two-tailed t-test: P=0.73). This result supports the hypothesis that β-parvin knockdown suppressed muscle differentiation. Since β-parvin knockdown was incomplete, heterogeneous, and transient, it is likely that the lower contractile force in the siParvb micro-tissues after 5 days of differentiation reflects a lower percentage of differentiated muscle fibers that were nonetheless similarly mechanoadaptive as muscle fibers under control conditions.

## Discussion

In this study, we show that cardiac and skeletal muscle contractility adapts to environmental stiffness. As a model system, we use muscle micro-tissues that self-assemble between two elastic pillars. The environmental stiffness experienced by the muscle can be varied over three orders of magnitude by changing the effective spring constant of the pillars. We then estimate the contractile forces from the bending of the pillars. Our data show that the active contractile forces of cardiac and skeletal muscle micro-tissues strongly increase with the pillar spring constant according to a power-law relationship. The power-law exponent provides a quantitative measure of mechanoadaptation, whereby values around zero indicate absence of mechanoresponsiveness, and values near unity indicate a linear increase of force generation with environmental stiffness. Knock-down of β-parvin in skeletal muscle micro-tissues drastically reduces contractile forces but does not affect the power-law exponent. Therefore, mechanoadaptation appears to be an intrinsic property of muscle cells, regardless of absolute force generation in the muscle tissue.

Single-cell studies have recapitulated the correlation between external stiffness and contractility in both striated [26] and smooth muscle cells [27]. Micro-tissues allow *in vitro* studies to be performed in a more realistic microenvironment under similarly controllable laboratory conditions and with similar experimental throughput. Moreover, it is possible to precisely tune the mechanical boundary conditions by changing the effective spring constant of the flexible pillars between which the tissues develop. This approach can be used to systematically investigate the contractile forces of micro-tissues over a sufficiently wide range of external stiffness and with a sufficient number of samples for a statistically sound evaluation.

In a previous study using such an approach, an increase in active and static forces of neonatal rat cardiomyocyte micro-tissues was observed at an external spring constant of 0.45 µN/µm compared to a spring constant of 0.2 µN/µm [11]. However, the authors of this study did not systematically investigate this effect and suggested that the higher contractility in stiffer environments was a consequence of better tissue compaction. In our experiments, by contrast, we did not observe large differences in micro-tissue compaction at different environmental stiffnesses.

Another study presented a system in which the boundary conditions of cardiac micro-tissue derived from human induced pluripotent stem cells (iPSCs) are regulated by real-time feedback control such that the pillars are pulled apart simultaneously with tissue contraction [28]. Under such near-isometric boundary conditions, contractile forces increased twofold compared to the unregulated condition, demonstrating that mechanoadaptation can be a very rapid process. In addition, iPSC micro-tissues also show a long-term adaptation of contractile forces to environmental stiffness, as shown in a study where the tissues were exposed to pillar spring constants between 0.1 to 10 µN/µm [29]. In close agreement with our data, that study reported an approximately 4.5-fold increase in contractile forces (from 57 µN to 262 µN) with higher stiffness, and a more pronounced maturation of the myocytes, as evidenced by an increased ratio of myosin heavy chain-beta to myosin heavy chain-alpha.

Active forces are measurable in C2C12 micro-tissues only after approximately 3-4 days in culture, after which the force magnitude increases until day 7 in culture. This temporal force increase correlates with the degree of desmin expression and the emergence of a higher power-law exponent. The increasing expression levels of desmin as well as the appearance of active contractile forces after electric pacing after day 3-4 indicate an increasing degree of myoblast-to-myotube differentiation. Fully differentiated muscle displays a regular striation pattern of actin and myosin filaments, as seen for example in Fig. 4B, that exhibit very slow turn-over [30]. Accordingly, treatment with the actin polymerization inhibitor cyto-D had little or no effect on the magnitude of active forces of differentiated C2C12 micro-tissues, consistent with previous reports [31]. By contrast, static forces of cardiac and skeletal micro-tissues, which we attribute to cells that have not differentiated to myotubes, decrease to the level of permeabilized tissues within minutes after treatment with cyto-D. Taken together, in agreement with a previous report [29], our data suggest the mechanical adaptation to environmental stiffness is facilitated by a faster and higher degree of cell differentiation.

The observation of higher contractile forces in response to a higher environmental stiffness, however, could potentially be explained by two trivial mechanisms. First, muscle tissue shortening is limited by the maximum distance of actomyosin filament sliding within a sarcomere [24, 32, 33]. This limit will be reached more readily in the case of softer pillars, resulting in seemingly smaller maximum forces. This mechanism, however, cannot account for our findings as the deflections of stiffer pillars tended to be smaller than the deflection of softer pillars (Fig. 5D,E). As a second explanation, one could consider the pillar-micro-tissue system as an arrangement of three Hookean springs (2 x *k_pillar_* and *k_tissue_*) [34]. Accordingly, when *k_pillar_* and *k_tissue_* are similar, substantial portions of the contractile forces are carried by the tissue and are not observable by pillar bending. Hence, the pillar deflection reports the true contractile force only when *k_pillar_ >> k_tissue_* (Fig. 7F). Indeed, *k_pillar_* was considerably larger than *k_tissue_* for the vast majority of our measurements (Fig. 7D). Moreover, *k_tissue_* of C2C12 tissues remained nearly constant over several days, whereas the degree of mechanoadaptation increased greatly during the same time period. In addition, a fit of the "three-spring-model" to the *F* versus *k_pillar_* measurements cannot recapitulate the mechanoadaptation response and fails especially at higher spring constants (Fig. S2). Thus, we conclude that the observed mechanoadaptation responses of muscle tissues are a biological phenomenon and cannot be attributed to such trivial mechanisms.

Micro-tissues in which the focal adhesion protein *β*-parvin has been knocked down show reduced contractile forces and a lesser degree of differentiation as judged from the larger twitch time constant. Reduced or absent desmin expression levels in muscle tissue have previously been shown to be accompanied by reduced force generation [23, 35] and possibly impaired functional adaptive response to mechanical overload [36]. Our finding that muscle tissues with reduced force generating ability after *β*-parvin knock-down are similarly mechanoresponsive as control tissues is therefore surprising. A likely explanation is a heterogeneous knock-down of *β*-parvin such that a fraction of the cells differentiated normally and displayed unaltered mechanoresponsiveness. Accordingly, the mechanoadaptation is a cellular process, and large contractile forces at the tissue level are not required. This interpretation is furthermore supported by our finding that the degree of stiffness adaptation, as quantified by the power-law exponent, was highest after day 5-6 in culture when contractile forces had not yet peaked.

A limitation of our study is that the performance of muscle tissues grown from both primary cardiac cells and the skeletal muscle cell line C2C12 declines after about 7-9 days in culture. Therefore, we cannot determine if muscle tissues exposed to low environmental stiffness only differentiate more slowly but would eventually catch up in their maximum contractility if given sufficient time. This remains an open question and could be addressed by studying muscle tissues grown from induced pluripotent stem cells or from skeletal muscle stem cells (satellite cells), both of which have been shown to maintain contractility over a time span of at least several weeks.

In summary, our study demonstrates that developing skeletal and cardiac muscle tissue displays a surprisingly large degree of mechanoadaptation over a wide range of environmental stiffness conditions spanning nearly three orders in magnitude.

## Materials and Methods

### Isolation of neonatal rat cardiomyocytes

All experiments were performed in accordance with the guidelines from Directive 2010/63/EU of the European Parliament on the protection of animals used for scientific purposes. Organ harvesting and preparation of primary cell cultures was approved by the local Animal Ethics Committee in accordance with governmental and international regulations on animal experimentations (protocol: TS-9/16 Nephropatho).

Ventricular cardiomyocytes were isolated from hearts of neonatal rats (P3) as previously described [37]. In brief, upon decapitation, hearts were isolated and the atria removed. The remaining ventricles were minced and digested utilizing the Neonatal Heart Dissociation Kit and the gentleMACS Dissociator (both Miltenyi Biotech), according to the manufacturer’s instructions. To enrich cardiomyocytes, cells were preplated for 1.5 h in DMEM-F12/Glutamax TM-I (Life Technologies), supplemented with 10% fetal bovine serum (FBS, Biowest), 100 U/ml penicillin and 100 μg/ml streptomycin (Life Technologies). Non-attached cells were collected, pelleted (5 min at 300 g) and resuspended in DMEM-F12/Glutamax TM-I, supplemented with 3 mM sodium pyruvate (Sigma Aldrich), 0.5% Insulin-Transferrin-Selenium (100x, Life technologies), 0.2% bovine serum albumin (Sigma Aldrich) and penicillin/streptomycin.

### Cell culture

C2C12 cells were cultured at standard incubation conditions (37 °C, 5% CO_2_, 95% humidity) in Dulbecco’s modified Eagle medium supplemented with 15% fetal bovine serum (Sigma-Aldrich), 2% penicillin-streptomycin (Gibco), 2% L-glutamine (Gibco), 2% non-essential amino acid solution (Gibco), 1 mM sodium pyruvate (Gibco), and 60 mM HEPES buffer (Gibco).

### Fabrication and culture of micro-tissues

Micro-tissues were grown as described previously [23]. In short, we cast PDMS devices, each consisting of 18 small wells with dimensions of 3.6 × 1.8 × 2 mm (l × w ×h) with two pillars (diameter: 500 µm, height: 2.5 mm, center-to-center distance: 1.8 mm). Before cell seeding, the PDMS devices were incubated overnight with a solution of 1% pluronic acid F-127 (Sigma-Aldrich) in water to prevent cells and matrix proteins from adhering to the chamber walls (Fig. S4). Then, we aspirated the coating and pipetted a bottom layer of 6 µl ice-cold unpolymerized collagen solution into each well.

For micro-tissues from neonatal rat cardiomyocytes, 1 ml collagen solution consisted of 50 µl of 2 mg/ml collagen R (collagen I from rat tail veins, Matrix Bioscience), 48 µl from 4.2 mg/ml collagen G (collagen I from bovine calf skin, Matrix Bioscience), 12.4 µl NaHCO_3_ (23 mg/ml, Sigma-Aldrich), 12.4 µl 10×DMEM (Seraglob), 1.9 µl NaOH (1M), and 876 µl of dilution medium consisting of one volume part NaHCO_3_, one volume part 10×DMEM, and eight volume parts H_2_O, adjusted to pH 10 using NaOH.

To obtain contractile micro-tissues grown from C2C12 cells, it was necessary to increase the collagen concentration and to add Matrigel. Here, 1 ml of collagen solution consisted of 100 µl of 2 mg/ml collagen R, 95 µl of 4.2 mg/ml collagen G, 98 µl of 10.2 mg/ml Matrigel (Corning), 37.1 µl NaHCO_3_, 37.1 µl 10×DMEM, 3.7 µl NaOH (1M), and 629 µl of dilution medium.

After pipetting 6 µl of unpolymerized matrix solution for the bottom layer to the wells, we centrifuged the PDMS devices (1 min at 300 g) and incubated them under standard conditions for 1 h for polymerization. Then, we pipetted a top layer of 6 µl ice-cold unpolymerized matrix solution mixed with 5000 cells on top of each bottom layer and incubated the samples for another hour without prior centrifugation. Then we carefully pipetted 1 ml of culture medium on top of the wells in each PDMS devices and thereafter changed the medium daily.

Micro-tissues from neonatal rat cardiomyocytes were cultured in Dulbecco’s modified Eagle medium/Ham’s F-12 (Gibco) supplemented with 5% fetal bovine serum (Sigma-Aldrich), 5% horse serum (Gibco), 2% penicillin-streptomycin (Gibco), 2% L-glutamine (Gibco), and 20 µM Cytosin-1-β-D-arabinofuranosid (Sigma-Aldrich). Micro-tissues from C2C12 cells were cultured in C2C12 cell culture medium (see section Cell culture) for the first 24 hours after seeding. Afterwards, we switched to a differentiation medium consisting of Dulbecco’s modified Eagle medium supplemented 0.5% horse serum (Gibco), 0.5% Insulin-Transferrin-Selen (Gibco), 2% penicillin-streptomycin (Gibco), 2% L-glutamine (Gibco), 2% non-essential amino acid solution (Gibco), and 1mM sodium pyruvate (Gibco).

### Neural network detection of pillar deflection

We trained a neural network to detect the position of the PDMS pillars in the recorded bright-field images. We used a modified version of the OR-UNet [38] without batch normalization and a low number of filters (16, 32, 64, 128, 128, 128, 128 for the layers). We first pretrained the neural network for 200 epochs with a batch size of 15 on 965 images on which the circular pillar outline was manually labeled. Images were cropped to obtain an equal number of background and pillar pixels. After pre-training, we trained the neural network again on the whole images with a batch size of 1 for 200 epochs. Data was augmented during training with i) x-y flipping, ii) up to 2.5 degree rotation, iii) contrast degeneration, iv) Gaussian blur with a width of maximal 5 pixels, v) brightness adjustment, and vi) slight zoom (between 0.95x and 1.05x).

To detect the pillar positions, the images were passed through the network, thresholded, topologically “closed” (binary dilation followed by erosion), and analyzed using the function *regionprops* from the Python scikit-image library to determine the x-y position of areas larger than 500 pixels. The automatic evaluation determined the distances between pillars with only a 0.4% relative standard deviation from the manually labeled distances. Automatically determined pillar centers were on average 3 pixels away from the hand labeled pillar centers.

### Analysis of contractile force

The axial contractile force *F* of the micro-tissues was determined from the pillar deflections *δ* as described previously [23]. In short, the relationship of *F* and *δ* is given by Hooke’s law

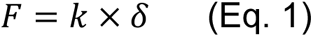

where *k* is the pillar’s spring constant at the height of the tissue above the pillar base *h*. *k* is computed from the the Euler-Bernoulli beam equation

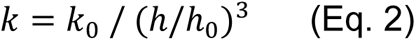

where *h_0_* is an arbitrarily chosen reference height of 1 mm and *k_0_* is a characteristic spring constant at that height. The value of *k_0_* depends on the PDMS’s crosslinker-to-base ratio, curing time, and curing temperature. We determined *k_0_* for PDMS pillars of all curing conditions used for this study from the force-bending relationship using a micromanipulator (Eppendorf InjectMan NI2).

We distinguish between static and active forces of micro-tissues. Static force is defined as the force resulting from pillar bending of myocytes that are not electrically paced.

Active force is defined as the contractile force in response to electrical pacing above the static force. Active force is calculated from the maximum pillar deflection during contraction compared to the length of the micro-tissue in the unpaced state. Bipolar electrical field stimulation was applied using a custom-made pacing device with four electrodes as described previously [23].

### Immunofluorescence staining

Cardiac and skeletal muscle micro-tissues were fixed and stained within the PDMS devices. Fixation was performed using 4% paraformaldehyde in phosphate-buffered saline (PBS) for 20 min, followed by three times washing with PBS for 10 minutes. The micro-tissues were then permeabilized in 0.5% Triton X-100 in PBS for 5 min and blocked in 3% bovine serum albumin (BSA) in PBS before they were incubated with primary antibodies (anti-sarcomeric alpha actinin (Abcam #9465) or desmin (Cell Signaling #5332)) at a dilution of 1:250 in blocking solution at 4 °C overnight. The next day, the samples were washed three times in PBS for 10 minutes. The micro-tissues were then incubated with a secondary antibody (either Jackson #715-025-150 or #711-095-152) at a dilution of 1:200 in blocking solution for 2 h, and then washed three times in PBS for 10 minutes. For the first washing step, the nuclear dye DRAQ5 (Abcam) was added to the PBS at a concentration of 1.7 µM.

### Imaging

Bright-field images were taken with a complementary metal-oxide semiconductor camera (Basler acA4112-30um). Confocal images were taken with an upright SP5X laser scanning microscope (Leica) using a 20X/1.0NA dip-in water-immersion objective lens. We recorded image stacks of 755 × 755 µm at a z-section distance of 2 µm for a depth of 150 µm into the micro-tissues. We maximum-projected the confocal image stacks of different wavelength channels and merged them using the image processing software ImageJ.

### Statistical methods

We used a two-tailed Student’s *t* test to compare groups. A *P* value < 0.05 was considered significant. In scatter plots, we examined the correlation between x-values and y-values by means of the coefficient of correlation (*R^2^*) and the Pearson correlation coefficient (ρ) as indicated in the figures. In bar graphs, whiskers represent standard errors. In boxplots, median values are represented by lines, and whiskers represent 5% and 95% percentiles.

### Force-tissue stiffness relationship in the absence of mechanoadaptation

To predict how tissue stiffness influences the relationship between contractile force and environmental stiffness when the tissue is not mechanoadaptive, we consider that the two pillars and the tissue (Fig. 7B) can be thought as being connected to walls at both ends (Fig. 7E). The tissue-generated active or static force *F*, which we in the following assume to be constant (i.,e. not mechanoadaptive) acts on the two pillars and is directed toward the center of the micro-tissue. As a result, the center of the micro-tissue remains stationary during contraction. This is equivalent to the situation where *F* acts on a parallel arrangement of a pillar and half of the micro-tissue connected to a wall on one side, which doubles the effective spring constant of the tissue so that the total spring constant of the arrangement is *k = k_pillar_* + *2×k_tissue_*. We assume that the tissue behaves as a linear Hookean spring with the same spring constant in extension and compression so that *F* = (*k_pillar_* + 2*×*k_tissue_) *× δ_pillar_*. To visualize how the measured force from the pillar deflection deviates from the tissue-generated force as a function of tissue stiffness, we plotted the ratio of *F_pillar_* / *F* against the ratio of *k_pillar_* / *k_tissue_* (Fig. 7F).

### Time course of a contractile force twitch

The speed of force build-up and relaxation during a single twitch depends chiefly on the acto-myosin bridge dynamics and the rate of Ca^2+^ ion reuptake by the sarcoplasmic reticulum. The Ca^2+^ ion reuptake rate distinguishes low and fast twitch muscle fibers and reflects the degree of muscle differentiation independent of the number of differentiated fibers in a tissue and the total active force [39, 40]. Ca^2+^ ion release into the cytoplasm can be regarded as near-instantaneous after delivering an electric pulse, *i.e.* it can be modeled as a delta-Dirac pulse, and the reuptake approximately follows a first-order kinetic process, hence it is well-described by a single exponential function. The time course of contractile force in turn follows the cytoplasmic Ca^2+^ concentration also according to a first-order kinetic process so that it can be modeled as a Dirac-pulse that is low-pass filtered twice [41]. For simplicity, we chose the time constants of the two low-pass filters to be equal so that we fit only a single time constant to the force versus time signal, normalized to the maximum force after the static force has been subtracted.

Despite having only one free fit parameter, the quality of the fit is excellent (Fig. 8E), and we can thus regard the fitted time constant as a measure for the rate of force buildup and force relaxation.

## Acknowledgments

We thank Igor Adameyko for providing C2C12 cells, Jana Petzold and Robert Becker for isolating cardiac muscle cells, and Christoph Schmidt for helpful discussions.

## Funding

- B.F. acknowledges support from the German Research Foundation (DFG) grants FA-336/12.1, FR-2993/23.1, HA3309/3-1, HA3309/6-1, HA3309/7-1, TRR 225 project 326998133 (subprojects A01, B08, and C02).
- F.B.E. acknowledges support from the DFG; TRR 225 project 326998133 (subproject C01).

## Author contributions

- D.K., J.L., T.W., and D.B. performed the experiments.
- D.K., M.S., C.A.D., J.K., S.W., I.T., and B.F. conceived and developed the micro-tissues assay.
- D.K. and B.F. analyzed the data.
- D.K., S.L.V., and I.T. conceived the genetic experiments.
- S.L.V., T.U.E., F.B.E., and I.T. provided material.
- D.K., R.C.G., and B.F. conceived and programmed software for data evaluation.
- D.K. and B.F. wrote the manuscript.
- D.K. and D.B. created the figures.
- All authors discussed the results and contributed to the final manuscript.

## Competing interests

The authors declare that they have no competing interests.

## Data and materials availability

All data needed to evaluate the conclusions in the paper are present in the paper and/or the Supplementary Materials. Raw data and additional data related to this paper can be provided from the corresponding author on request.

If applicable, begin the section with text that acknowledges non-author contributions. (Note this section does not have a general heading).

## Animal ethics statement

All animal experiments (harvesting neonatal rat hearts to isolate cardiomyocytes) were performed in accordance with the guidelines from Directive 2010/63/EU of the European Parliament on the protection of animals used for scientific purposes. In addition, all experiments were approved by the local Animal Ethics Committee in accordance with governmental and international regulations on animal experimentations (protocol: TS-9/16 Nephropatho).

---

**Fig. S1.**
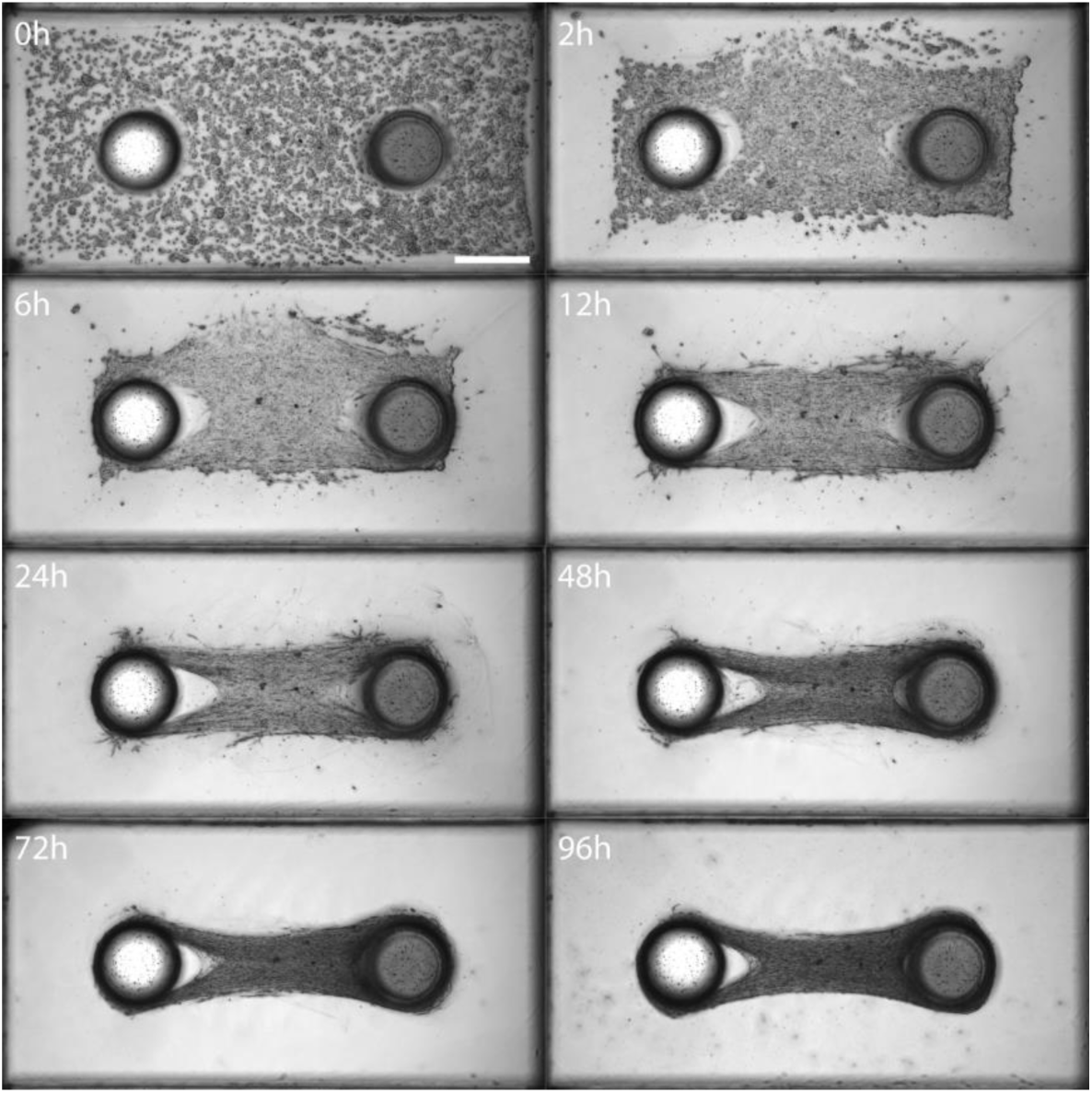
Compaction of cells and matrix into micro-tissues. Minimum intensity projections of bright-field images of a micro-tissue at different times after cell seeding. Upon polymerization, C2C12 cells adhere to the collagen-I/Matrigel matrix, exert forces and subsequently cause collective remodeling of the network. Scale bar: 500 µm.

**Fig. S2.**
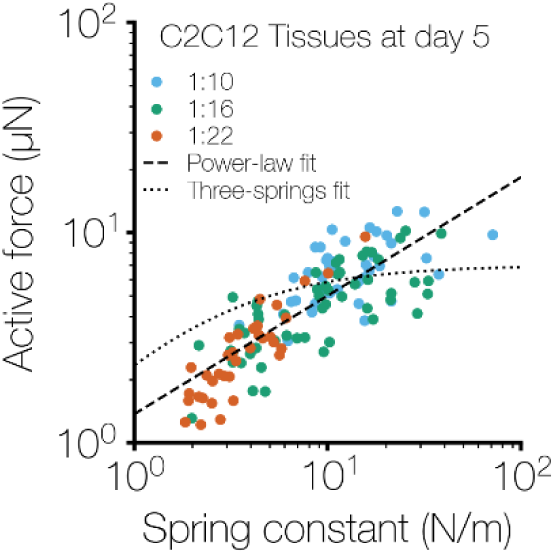
Power-law fit of active forces of C2C12 micro-tissue compared to a three-spring-model. The same scatter plot as in Fig.5B for day 5 (*i.e.* the day of weakest mechanoadaptation). The dashed line represents a power-law fit as before and the dotted line represents a fit based on a three-spring-model optimized by least-squares error minimization.

**Fig. S3.**
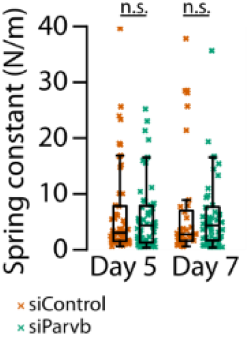
Spring constants of PDMS pillars for experiments with control- and Parvb-siRNA-treated C2C12 micro-tissues are not significantly different. Two-tailed t-test; n.s., P≥0.05.

**Fig. S4.**
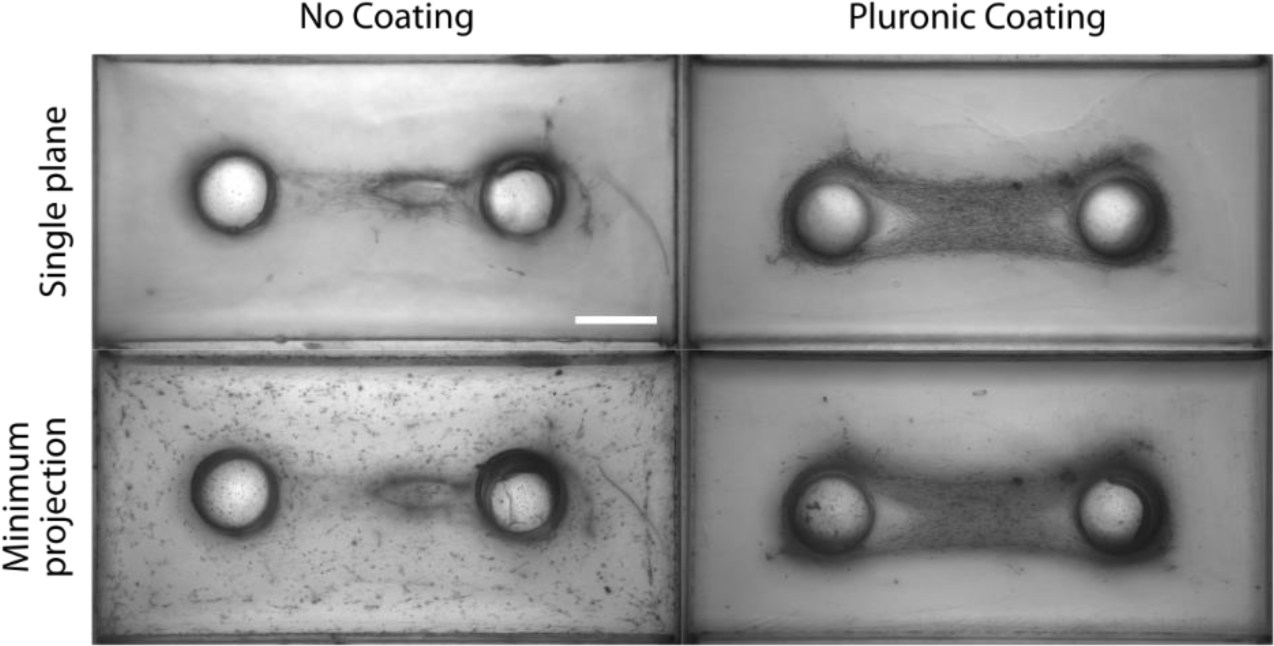
Pluronic acid coating prevents matrix adhesion to the PDMS chamber. Bright-field images focused on two micro-tissues 24 hours after cell seeding (top), and minimum projections of bright-field images of the same samples (bottom). The PDMS device in the left column was not coated with pluronic acid prior to micro-tissue fabrication, while the PDMS device in the right column was. Compared to the coated sample, the micro-tissue in the uncoated PDMS device did not form properly because cells and matrix adhered to the walls of the chamber, especially at the bottom. Scale bar: 500 µm.

## Notes

### Competing Interest Statement

The authors have declared no competing interest.

